# Effects of exposure to sublethal concentrations of methoxyfenozide on honey bee colony activity and thermoregulation

**DOI:** 10.1101/416354

**Authors:** William G. Meikle, Vanessa Corby-Harris, Mark J. Carroll, Milagra Weiss, Lucy A. Snyder, Charlotte A.D. Meador, Eli Beren, Nicholas Brown

## Abstract

Methoxyfenozide is an insect growth regulator (IGR) commonly used in agricultural to simultaneously control pests and preserve beneficial insect populations; however, its impact on honey bees in not fully understood. We conducted field and laboratory experiments to investigate bee health in response to field-relevant doses of this pesticide. Significant effects were observed in honey bee colony flight activity and thermoregulation after being treated with methoxyfenozide. Data collected indicated that hives fed 500 ppb methoxyfenozide treated pollen patty had: 1) a significantly reduced rate of daily hive weight loss due to forager departure at the start of the colony’s daily activity; 2) the end of the colony’s daily activity delayed by 17-21 minutes compared to Control; and 3) higher temperature variability during the winter. Colonies in the 125 ppb treatment group had fewer differences with the Control group, but did show a delay in the foraging end time by 30-46 minutes compared to the Control. Bee colony metrics of adult bee mass and brood surface area, and individual bee measurements of head weight, newly-emerged bee weight, and hypopharyngeal gland size were not significantly affected by the methoxyfenozide exposure levels of our experiments. An experiment conducted using the same treatment groups in the spring resulted in fewer differences among groups than did the experiments conducted in the fall. Analyses of methoxyfenozide concentrations in the treatment patty, wax, and bee bread showed that: 1) observed methoxyfenozide concentrations were about 18-60% lower than the calculated concentrations; 2) no residues were observed in wax in any treatment; and 3) methoxyfenozide was detected in stored bee bread in the 500 ppb treatment, at concentrations about 1-2.5% of the observed concentration for that treatment. These results suggest that there may be significant effects on honey bee colony behavior (and possibly health) in the field that are difficult to detect through traditional hive inspections and individual metrics.

## Introduction

Honey bee colonies are frequently exposed to agrochemicals, including many different classes of insecticides, among them insect growth regulators (IGRs) [1]. Broadly speaking, IGRs interfere with insect growth and development in their target pest species [2]. The IGR methoxyfenozide is an ecdysone receptor agonist that binds the ecdysteroid receptor and activates the ecdysteroid signaling pathway [3, 4]. Unlike the ecdysteroid hormone that precisely controls larval development by binding to the receptor for a defined period of time, ecdysteroid agonists bind irreversibly, disrupting the expression of genes involved in cuticle development, sclerotization, and ecdysis [4,5]. Methoxyfenozide specifically targets lepidopterans and research shows that it has high affinity for the lepidopteran ecdysteroid receptor that is not seen in other insect orders (reviewed in [3, 4]). Although the binding specificity of methoxyfenozide to the *Apis mellifera* ecdysteroid receptor has not been determined, it is safe to assume that it should not bind. Fewer non-target effects makes methoxyfenozide an attractive, targeted solution for controlling pests while preserving beneficial insect populations. Methoxyfenozide is registered in more than 50 countries for use in a variety of crops, including those pollinated by honey bees. Methoxyfenozide use has increased 15-fold between 2001 and 2015, from ∼30,000 to ∼450,000 pounds annually [6], mostly in orchards, to control lepidopteran pests like the navel orangeworm, *Amyelois transitella* (Walker) [7].

Reported toxicity of methoxyfenozide to young worker bees is low, with an acute toxicity LD_50_ greater than 100 ug/bee [8] and, when formulated with spinetoram, an oral toxicity LC_50_ of about 712 mg/L (ppm) in young workers [9]. Nevertheless growers have been advised to avoid spraying during bloom because the impact of methoxyfenozide on earlier life stages, older workers, or on colonies is not fully understood [10]. Despite these spray recommendations, methoxyfenozide has been detected in commercial colonies in bees (9-21 ppb) and hive materials including wax (80-495 ppb), pollen (35-128 ppb), and honey (3 ppb) [1, 11]. In a recent study, methoxyfenozide has been shown to decrease forager survival at field-relevant doses [12].

Other IGRs have caused delayed sublethal effects in adult honey bees after larval exposure. Larval bees exposed to the juvenile hormone analog pyriproxyfen in pollen patty at 321 ppb showed increased deformities as adults, reduced adult survivorship and exposed adults had difficulties integrating into the general adult population; fewer effects were observed at the lower concentration of 129 ppb [13]. The impact of methoxyfenozide on honey bee colonies is not fully understood. Although methoxyfenozide is marketed as a safe pesticide due to its specificity to certain pests, it is nonetheless possible that honey bees are negatively impacted simply due to the energetic cost of detoxifying an exogenous chemical [14-16]. Pesticide exposure can lead to increased expression of stress response and detoxification genes [17], but expression patterns vary [18]. It is unclear whether detoxification, *per se*, is stressful to honey bees, although metabolomic analyses of bees exposed to plant secondary toxins suggest that it is [14] and nurse-aged bees exposed to some classes of pesticides have reduced hypopharyngeal glands [18, 19], a pattern observed with other stressors [20].

Effects that are difficult to detect on the level of the individual may be detected on the level of the colony [21]. Continuously monitoring weight and internal hive temperature of honey bee hives has provided information on bee colony growth and activity [22-24]. In those studies, continuous weight and hive temperature data were detrended by subtracting the 25-hour running average from the raw data. Running average weight data provided information on longer-term colony growth while the detrended within-day data were modeled using sine curves to yield information on foraging activity and success. Similarly, the average and detrended hive temperature data were related to capped brood levels [22]. These approaches were used to detect colony-level treatment effects of sublethal exposure of the neonicotinoid pesticide imidacloprid on flight activity and internal hive temperature control (colony thermoregulation) [25]. In another approach, continuous hive weight data were detrended by subtracting the value at midnight, rather than the running average, from each of the values of that day until the next midnight [26]. In that approach, a piecewise regression model was fitted to single-day datasets in the original time scale using an R function [27] based on a bootstrapping method [28]. Several parameters of the piecewise regression provide biological or behavioral interpretations, such as break points around dawn and dusk reflecting the beginning and ending of the daily active periods for the hive [26].

To more fully address the possible consequences of methoxyfenozide exposure on honey bees, we conducted replicated field and laboratory investigations of bee health in response to field-relevant doses of this pesticide. In two different years, we mixed methoxyfenozide into pollen patty and fed it to colonies. Colony parameters such as brood area, adult weight, foraging activity, and hive temperature were measured. We find that methoxyfenozide does not exert a massively negative effect on honey bees, but that small differences among hives in foraging activity and hive temperature regulation appear to be exacerbated in the high-dose treatments compared to the control.

## Materials and methods

### 1. Preparation of pollen supplement

A 50 mg/ml stock solution of methoxyfenozide was prepared in acetone. 18 kg of pollen patty supplement was prepared in 1.5 kg batches in a stand mixer at a ratio of 1:1:1 corbicular pollen (Great Lakes Bee Co.):granulated sugar:drivert sugar (Domino Foods). For the 500 ppb treatment, 15 µL of the stock solution of methoxyfenozide was added to 235 µL of acetone yielding a 3 mg/ml solution. 250 µL of this 3 mg/ml solution was added to 130 ml of water and then thoroughly mixed with 1.5 kg of patty. Similarly, the 125 ppb treatment patty was prepared by adding 3.75 ul of the stock solution to 246.25 ul acetone yielding a 0.75 mg/ml solution. 250 µL of this 0.75 mg/ml solution was added to 130 mL water and thoroughly mixed into 1.5 kg of patty. All of the diet was prepared at the same time for the Fall 2016 experiment. The diet was divided into 100 g patties and stored at -20°C until the supplement was fed.

Pollen supplement for Fall 2017 and Spring 2018 experiments was prepared in a similar manner except that the methoxyfenozide stock solution was prepared to 6 mg/ml in acetone. 125 µL of that solution was then mixed with 100 mL water for a concentration of 0.75 mg/100 mL and applied to 1.5 kg supplement for a concentration of 500 ppb. Likewise, 31 µL of stock solution was diluted with 93 µL acetone and mixed with 100 mL water per 1.5 kg diet for a concentration of 125 ppb. Control patties were treated with 125 µL pure acetone and 100 mL water. Patties were divided and stored in the same manner. Patty samples (3 g) were submitted to the Laboratory Approval and Testing Division, Agricultural Marketing Service, USDA (LATD), Gastonia, NC, to determine methoxyfenozide concentrations.

### 2. Fall 2016 field experiment

In August 2016, eighteen honey bee colonies were selected from apiaries at or near the Carl Hayden Bee Research Laboratory, USDA-ARS, Tucson, AZ (32°16‘30.17“N, 110°56‘28.52“W) and the Santa Rita Experimental Range (SRER) (31°46‘38.08“N, 110°51‘47.39“W) and moved to a single site at SRER in August, 2016. Colonies had been stocked with Cordovan-Italian queens (C.F. Koehnen & Sons, Glenn, CA) and housed in painted, 10-frame wooden Langstroth deep boxes fitted with migratory wooden lids (Mann Lake Ltd, Hackensack, MN). Colonies were 6-18 months old at the start of the experiment. The SRER apiary was provided with a permanent water source and hives were spaced 1-3 m apart. Hives were placed on stainless steel electronic scales (TEKFA^®^ model B-2418 and Avery Weigh-Tronix model BSAO1824-200) (max. capacity 100 kg) connected to 12-bit dataloggers (Hobo^®^ U-12, Onset Computer Corporation) that were set to record weight every 5 minutes. The system had an overall precision of approximately ±20 g. On the same day, a hive temperature sensor (iButton Thermochron, precision ±0.06°C) enclosed in plastic tissue embedding cassettes (Thermo Fisher Scientific, Waltham, MA) was stapled to the center of the top bar on the 5th frame in each hive and set to record every 30 min.

On 29-30 August, 2016, hives were given a full pre-treatment assessment (see [24, 29]). Briefly, the hive was opened after the application of smoke, and each frame was lifted out, gently shaken to dislodge adult bees, photographed using a 16.3 megapixel digital camera (Canon Rebel SL1, Canon USA, Inc., Melville, NY), weighed on a portable scale (model EC15, OHaus, 15 kg max. cap.), and replaced in the hive. Frames were removed and replaced sequentially. During this first assessment (but not subsequent assessments), all hive components (i.e. lid, inner cover, box, bottom board, frames, entrance reducer, internal feeder) were also shaken free of bees and weighed to yield an initial mass of all hive components. At the initial inspection, 3-5 g of wax were collected from each hive into 50 ml centrifuge tubes and stored at -80°C; samples collected in September, prior to treatment, were pooled and subjected to a full panel analysis for residues of 192 pesticides and fungicides, from all major classes, by LATD. Samples from later assessments were pooled within treatment group and subjected only to methoxyfenozide residue analysis.

The total weight of the adult bee population (whole colony adult bee mass) was calculated by subtracting the combined weights of hive components obtained in the pre-treatment assessment from the total weight of bees and hive materials recorded the midnight prior to the inspection. The area of sealed brood per frame was estimated from the photographs using ImageJ version 1.47 software (W. Rasband, National Institutes of Health, USA). The 18 colonies were randomly assigned to three treatment groups, to be given the Low (125 ppb), High (500 ppb) and Control treatments. Care was taken to ensure the average adult bee mass per treatment group varied by no more than 100 g among all treatments to minimize pre-existing differences (Table 1). Each hive was given 100g pollen patty twice a week with the appropriate concentration starting on 2 September, continuing weekly for 9 weeks until 28 October (1800 g treatment pollen patty total per hive). Consumption of the patty was measured by weighing any patty (wet weight) that remained after one week. Hives were assessed on 2 November, 2016 (first post-treatment assessment, 61 d after initial treatment). Each colony was then given 3 kg of 1:1 sugar syrup and frame of capped honey on 10 November because of low food stores. Colonies were assessed and sampled for the final time on 30 January, 2017, (2^nd^ post-treatment assessment) to determine the long-term effects of methoxyfenozide exposure on overwintering.

**Table 1.** Adult bee masses and brood surface areas for the Fall 2016, Fall 2017 and Spring 2018 field experiments.

| Dates | Adult bee mass (g) |  |  | Capped brood surface area (cm<sup>2</sup>) |  |  |
| --- | --- | --- | --- | --- | --- | --- |
|  | 125 ppb | 500 ppb | control | 125 ppb | 500 ppb | control |
| Fall 2016: |  |  |  |  |  |  |
| 30 Aug. 2016 | 2166 ±141 | 2254 ±234 | 2202 ±204 | 2120.6 ±220.7 | 1972.4 ±436.0 | 2475.5 ±291.7 |
| 2 Nov. 2016 | 1550 ±153 | 1498 ±330 | 1633 ±139 | 583.2 ±53.1 | 498.0 ±134.6 | 546.6 ±97.7 |
| 1 Feb. 2017 | 782 ±224 | 777 ±296 | 1352 ±107 | 9.1 ±8.0 | 3.7 ±0.6 | 11.6 ±7.9 |
| Fall 2017: |  |  |  |  |  |  |
| 13 Sep. 2017 | 1641 ±251 | 1591 ±208 | 1595 ±141 | 426.4 ±112.5 | 630.1 ±158.3 | 756.7 ±98.3 |
| 15 Nov. 2017 | 1563 ±170 | 1496 ±224 | 1434 ±83 | 274.9 ±38.4 | 210.6 ±33.1 | 269.4 ±53.0 |
| 13 Feb. 2018 | 1010 ±230 | 920 ±150 | 810 ±90 | 412.8 ±96.2 | 320.1 ±125.4 | 496.7 ±153.6 |
| Spring 2018: |  |  |  |  |  |  |
| 18 Apr. 2018 | 1211 ±422 | 1453 ±416 | 780 ±136 | 1607.2 ±413.6 | 2613.8 ±640.1 | 1055.6 ±245.7 |
| 23 May 2018 | 1054 ±351 | 1553 ±491 | 534 ±162 | 1110.6 ±169.9 | 1292.7 ±259.2 | 477.3 ±167.7 |

### 3. Fall 2017 field experiment

The experiment described above was repeated in 2017. In April, 2017 twenty colonies were started from packages from the same supplier as the previous year and installed in SRER at a site about 2 km away from the Fall 2016 site. Colonies were in single deep hive boxes and maintained with new Cordovan-Italian queens from the same queen supplier as the first year. Colonies were fed supplemental pollen patty in the spring, and 12 kg 1:1 sugar syrup between May and early September. On 13 September 2017, full pre-treatment hive assessments were conducted on all colonies. Second deep boxes were added to 6 of the colonies because of their size. Colonies were divided into 3 groups with similar adult bee masses (<100 g difference), and each colony was given 100 g treatment pollen patty twice per week beginning on 22 September and continuing for 8 weeks until 9 November (48 d after initial treatment, 1600 g treatment pollen patty total). Three supplemental feedings of 3 kg of 1:1 sugar syrup were provided to each colony during the treatment period. The 1^st^ post-treatment hive assessments were conducted on 15 November. Smaller colonies were reduced to single boxes for overwintering on 16 November 2017. The 2^nd^ post-treatment hive assessments were conducted on 13 February 2018. From 1 to 2 g of bee bread was collected from each hive at each assessment. As with the wax samples for the Fall 2016 experiment, samples collected in September, prior to treatment, were pooled and subjected to a full panel of residue analyses while samples from later assessments were pooled within treatment group and subjected only to methoxyfenozide residue analysis. Samples of protein patty from each treatment were also analyzed for methoxyfenozide concentration. In addition, newly-emerged bees (NEBs) were also sampled by pressing an 8 cm x 8 cm x 2 cm mesh queen cage into a section of capped brood, then returning the following day to collect NEBs that had emerged within the cage over the previous 24 h. The NEBs were then placed in a 50 mL centrifuge tube, frozen on dry ice, and stored at -80°C. At the laboratory, 5 bees per hive per assessment date were placed in Eppendorf tubes, weighed, dried for 72 h at 60°C, then re-weighed to determine average wet and dry weight per bee.

### 4. Spring 2018 field experiment

The hives used in the Fall 2017 experiment were retained in the same treatment groups and treated in the same manner as in the previous fall, using pollen patties with the same concentrations of methoxyfenozide. Treatments consisting of 100 g pollen patty were started on 8 March and continued twice weekly for 6 weeks until 13 April (42 d after initial treatment, 1200 g treatment pollen patty total). All hives were given 4 L supplemental sugar syrup on 30 March. Hives were evaluated on 18 April (1^st^ post-treatment assessment) and again on 24 May (2^nd^ post-treatment assessment). Bee bread and NEBs were sampled at each assessment.

### 5. Adult bee head weights from Fall 2017 experiment

Head weight was measured on samples of nurse-aged bees collected from all colonies after the November 2017 post-treatment hive evaluation. Nurse-aged bees visiting cells containing larvae for a period of ≥5 seconds were collected. The captured bees were immediately flash frozen in the field and were maintained at -80°C until their heads were weighed. To obtain head weights, each head was thawed and weighed using a using a Sartorius CP2P microbalance at a resolution of 0.01 mg (see [30]).

### 6. Hypopharyngeal gland size of nurse-aged workers fed methoxyfenozide

In order to assess whether oral methoxyfensozide exposure affected nurse-aged bee health, HPG size was measured in caged nurse-aged bees fed either 1000 ppb of methoxyfenozide in 30% sugar syrup or a control dose of methoxyfenozide-free sugar syrup. A comparatively high dose was chosen to increase the probability of a measurable effect. NEBs emerged overnight from brood frames taken from three colonies in a temperature-controlled dark room (32–34°C, 30–40% relative humidity). The next morning, the NEBs were distributed among eight cages to a density of 100 bees per cage. Cage dimensions were 11.5 x 7.5 x 16.5 cm, with narrow sides, top and base made of Plexiglas^®^ and the broad sides and floor made of 3mm aperture galvanized steel mesh. A plastic 50 mL bottle for distilled water and a plastic 30mL bottle for syrup, each with a small hole in the lid, were inverted and placed over holes on the top of each cage. Four cages were provided with 1000 ppb methoxyfenozide in 30% sucrose syrup and four cages with 30% sucrose syrup without the pesticide as controls. All cages were provided with pollen patty, 1:1:1 sucrose: drivert sugar : natural pollen (Great Lakes Pollen, Bulkfoods.com) *ad libitum*. The caged bees were maintained at 34°C and 30–40% relative humidity. At 8d after emergence, 10 bees per cage were flash frozen and maintained at -80°C until their HPGs were dissected and measured (see [31]). Between 10 and 12 HPG acini were measured for each gland to obtain an average HPG acinus size for each bee.

### 7. Data analysis: Hive assessment, patty consumption, NEB weights, nurse bee head weights and HPG size

Adult bee masses, capped brood surface areas and NEB dry weights were compared among treatments across sampling occasions using repeated-measures MANOVA (SAS version 9.4) with treatment, experiment and day as main factors, all 2-way interactions, and with pre-treatment adult bee mass as a covariate to control for pre-existing colony differences. Per colony patty consumption and nurse bee head weights were analyzed using ANOVA, in which treatment was the main effect and hive was a random effect. Caged-bee data were analyzed using ANOVA, with treatment as a fixed effect and cage replicate as a random effect. Post hoc contrasts with the Bonferroni correction for multiple comparisons were reported for significant treatment effects.

### 8. Data analysis: Hive weight

Continuous hive weight data were considered with respect to average daily weight and within-day changes. Weight data were detrended for each day by subtracting the hive weight estimate at midnight (or closest time thereafter) from each subsequent weight value over the next 24 h (see [25]). The resulting within-day weight datasets were modeled using the “segmented” function in R which fits a segmented line derived from a linear or generalized linear model to a dependent variable using a bootstrapping procedure [27]. Bee colonies outside of a nectar flow during winter tend to lose weight and exhibit consistent daily patterns (Fig 1). Piecewise regressions with 4 breakpoints were fit to the data, which yielded estimates for 10 parameters: 4 break point values, 5 slope values and the adjusted r^2^. Because the data were detrended by subtracting the raw data value at midnight, daily datasets were mathematically independent. A repeated measures MANOVA was conducted on these daily parameter values of interest (letters refer to Fig. 1):

**Figure 1.**
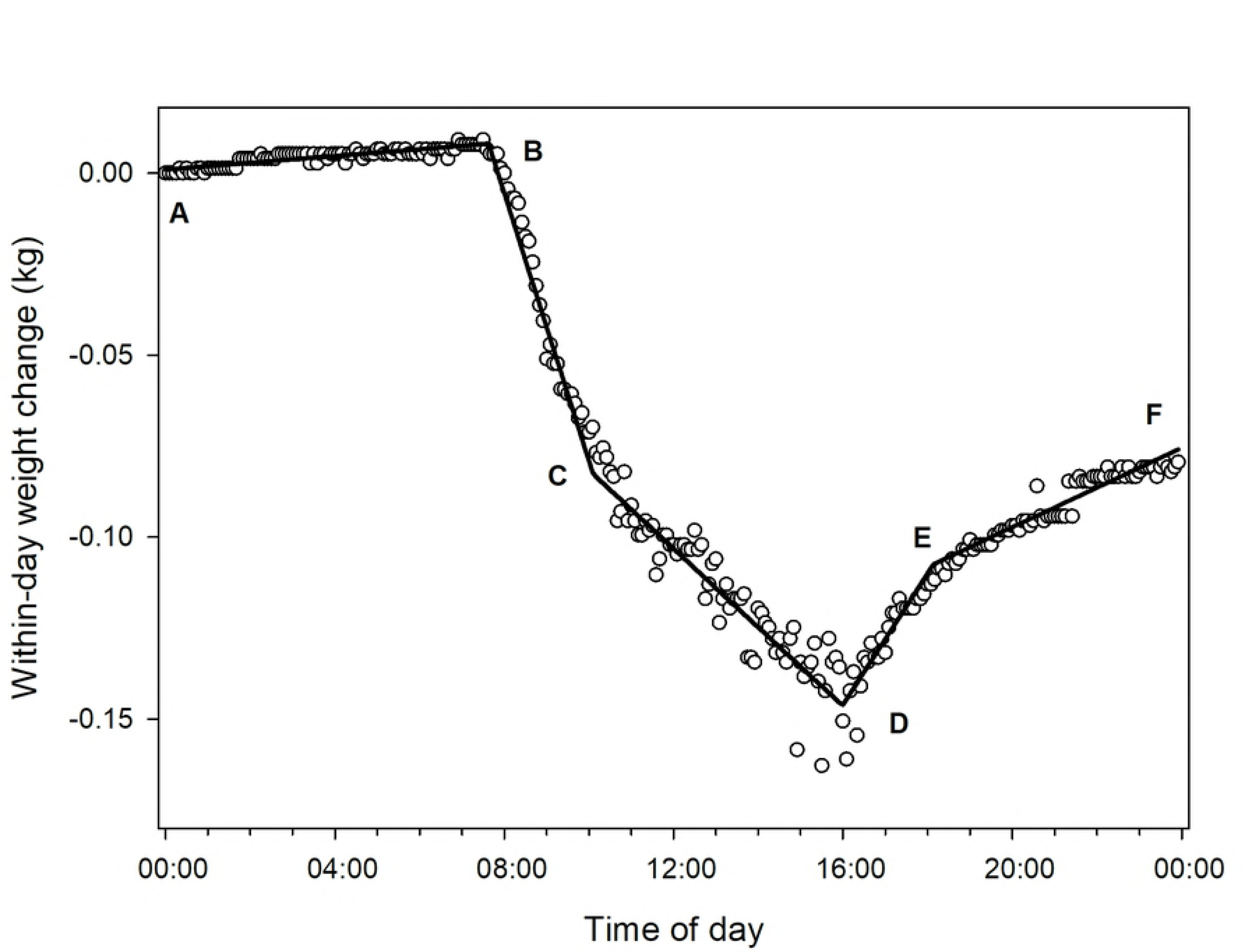
Graph showing data and fitted piecewise regression to data from a single hive in the Control group for a single day (24 November 2017) during a nectar and pollen dearth.

1. Beginning of initial forager departure (usually the 1^st^ break point, Point B);
2. End of initial forager departure (usually 2^nd^ break point, Point C);
3. Time of final forager return (usually the 4^th^ break point, Point E);
4. Average slopes of the 1st and 5th segments, which are weight changes at night when colonies are not foraging (Segments AB and EF);
5. Slope of the first segment after initial forager departure, usually the 2nd segment (usually Segment BC); and
6. Slope of the last segment before dusk, usually the 4th segment (usually Segment DE).

For statistical analysis, if the 1st break point occurred before 4AM, the 2nd break point was used as the time of initial forager departure (with no restrictions placed on that second estimate) and the slope of the 3rd, rather than 2nd, segment was used as the rate of weight loss due to forager departure. Likewise, if the 4th break point occurred after 8PM then the 3rd break point was taken as the time of final forager return (with no restrictions placed on that second estimate). For all analyses involving slopes, the pre-treatment adult bee mass was included as a covariate to control for pre-existing differences in forager populations.

1. **Point A:** First weight measure at midnight or shortly thereafter;
2. **Segment AB:** Inactive period in the early morning; hive weight change is likely due to bee respiration and changes in the moisture content of nectar, pollen and wooden hive parts;
3. **Point B:** Bee departure at beginning of active period (usually close to dawn);
4. **Segment BC:** Active period usually showing hive weight loss due to far greater numbers of departing bees compared to returning bees;
5. **Point C:** Point at which mass of returning bees increases relative to mass of departing bees;
6. **Segment CD:** Continued increased return of foragers;
7. **Point D:** Point at which mass of returning foragers, including nectar and pollen as well as bee weight loss due to respiration, exceeds the mass of departing bees plus weight loss due to drying and respiration in the colony;
8. **Segment DE:** Inactive period with hive weight change driven mainly by respiration and changes in ambient humidity – usually close to parallel with segment AB;
9. **Point E:** Return of bees to the hive around dusk is completed;
10. **Segment EF:** Inactive period in the late evening; similar dynamics with moisture content of hive, bee respiration and ambient humidity;
11. **Point F:** Last weight measure just before midnight.

### 9. Data analysis: Hive temperature

Internal hive temperature data were divided into daily average values and within-day detrended data. Detrended data were calculated as the difference between the 25 hour running average and the raw data [24]. Sine curves were fit to 3-day subsamples of detrended data taken sequentially by day, and curve amplitudes, representing estimates of daily hive temperature variability, were used as response variables. For hives with two boxes, only temperature data from the lower (brood) box were analyzed. Repeated measures MANOVA (Proc Glimmix, SAS Inc. 2002) was used to evaluate the effects of treatment, day, and their interaction, with the pre-treatment whole colony adult bee mass as a covariate on both the average daily hive temperature and the amplitudes of the fit sine curves. Temperature amplitude datasets were reduced to one value per hive point every 3 d for repeated measures analysis to ensure no overlap between subsamples.

## Results

### 1. Hive assessment

Methoxyfenozide treatment did not have a measurable impact on either whole colony adult bee mass (P=0.73) or brood surface area (P=0.43) in the Fall 2016 or Fall 2017 experiments, nor were the two experiments different from each other with respect to these metrics (P=0.36 and P=0.26, respectively) (Table 1). Considered separately, neither adult bee mass nor brood surface area in the Spring 2018 experiment was affected by treatment (P=0.14 and P=0.37, respectively). During the Spring 2018 experiment, two colonies in the 125 ppb treatment group and one colony in the 500 ppb treatment group died, in all cases around 1 May. Pre-treatment adult bee mass was significantly correlated with adult bee mass at the 2^nd^ post treatment assessment for the Fall 2016, Fall 2017 and Spring 2018 field experiments (adjusted r^2^ = 0.27, 0.44 and 0.31, respectively). Two colonies in the 500 ppb treatment group and one in the 125 ppb treatment group died in the Fall 2016 experiment, and one colony in the 500 ppb treatment group and two in the 125 ppb treatment group died in the Spring 2018 experiment.

### 2. Patty consumption

Bee colonies in all groups consumed all the pollen feed in the Fall 2016 and Fall 2017 experiments. In the Spring 2018 experiment some colonies did not consume all 1100 g patty but average values (wet weight) among treatment groups were not significantly different when average adult bee mass during feeding period was used as a covariate (P=0.72): the 500 ppb group consumed 900±97 g, the 125 ppb group consumed 868±154 g, and the control group consumed 871±94 g. Total consumption was related to colony size: the average adult bee mass of the colonies that consumed all the patty was 1.61 ±0.28 kg while that for the colonies that did not was 0.74±0.08 kg, and among colonies that did not finish the patty, consumption was directly proportional to adult bee mass (F_1,10_=13.21, P=0.0046, adj. r^2^ =0.53).

### 3. NEB weights and nurse bee head weights

Dry weights of NEBs were not significantly different among treatments in the Fall 2017 experiment (P=0.45). Methoxyfenozide application did not influence head weight of bees collected in the Fall 2017 experiment. Average head weights (±s.e.) for bees in hives treated with the 500 ppb (12.51±0.37 mg), 125 ppb (12.29±0.31 mg) and control (12.32±0.19 mg) treatments did not differ significantly (P=1.0).

### 4. HPG size of caged bees

Oral exposure to methoxyfenozide during young adult development did not impact the hypopharyngeal gland sizes of nurse-aged workers (P=0.31). The average (±s.e.) acinus size of bees exposed to 1000 ppb methoxyfenozide in syrup was 0.021±0.007 mm^2^, while those fed the control treatment had glands that were 0.025±0.007 mm^2^.

### 5.Pesticide analyses

Wax samples taken during the Fall 2016 experiment were analyzed for methoxyfenozide residues and none were found (Limit of Detection [LOD]=1 ppb) with the exception of trace amounts detected in the 500 ppb treatment in early November, just after the end of the treatment period. For the Fall 2017 experiment, bee bread samples were analyzed rather than wax. In the initial sample, analyzed with respect to a full panel of 192 compounds, only trace amounts of diphenylamine (LOD=2 ppb) and 118 ppb of thymol were detected. No methoxyfenozide was detected in the bee bread except for samples collected from hives in the 500 ppb treatment in the November, February and April hive assessments. Those samples contained 5, 2 and 2 ppb methoxyfenozide, respectively. The protein patty samples for the 500 ppb, 125 ppb and control treatments were found to have 199, 103 and 0 ppb methoxyfenozide, respectively.

### 6. Hive weight for Fall 2016 and Fall 2017 experiments

Piecewise regression curves fit the data well on average: average (±s.e.) adj. r^2^ values for the 125 ppb, 500 ppb and Control groups were 0.94 ±0.03, 0.92±0.05 and 0.91±0.05, respectively, for the Fall 2016 experiment and 0.96±0.02, 0.93±0.04 and 0.94±0.03, respectively, for the Fall 2017 experiment. Dusk break point was significantly affected by treatment (Fig 2). Post hoc contrasts showed that all treatment groups were significantly different from each other. Lower values in the Control group indicate that the dusk break point occurred significantly earlier in the day than for either of the other treatment groups, and earlier for the 500 ppb group than for the 125 ppb group. The two fall field experiments were themselves significantly different with respect to dusk break point. In the Fall 2016 experiment, the Control treatment group average dusk break point was about 4:37 PM, with the 500 ppb treatment group 21 minutes later and the 125 ppb treatment group 30 minutes later than Control. In the Fall 2017 experiment, the Control treatment group average dusk break point was about 4:59 PM, with the 500 ppb treatment group 17 minutes later and the 125 treatment group 46 minutes later than Control.

**Figure 2.**
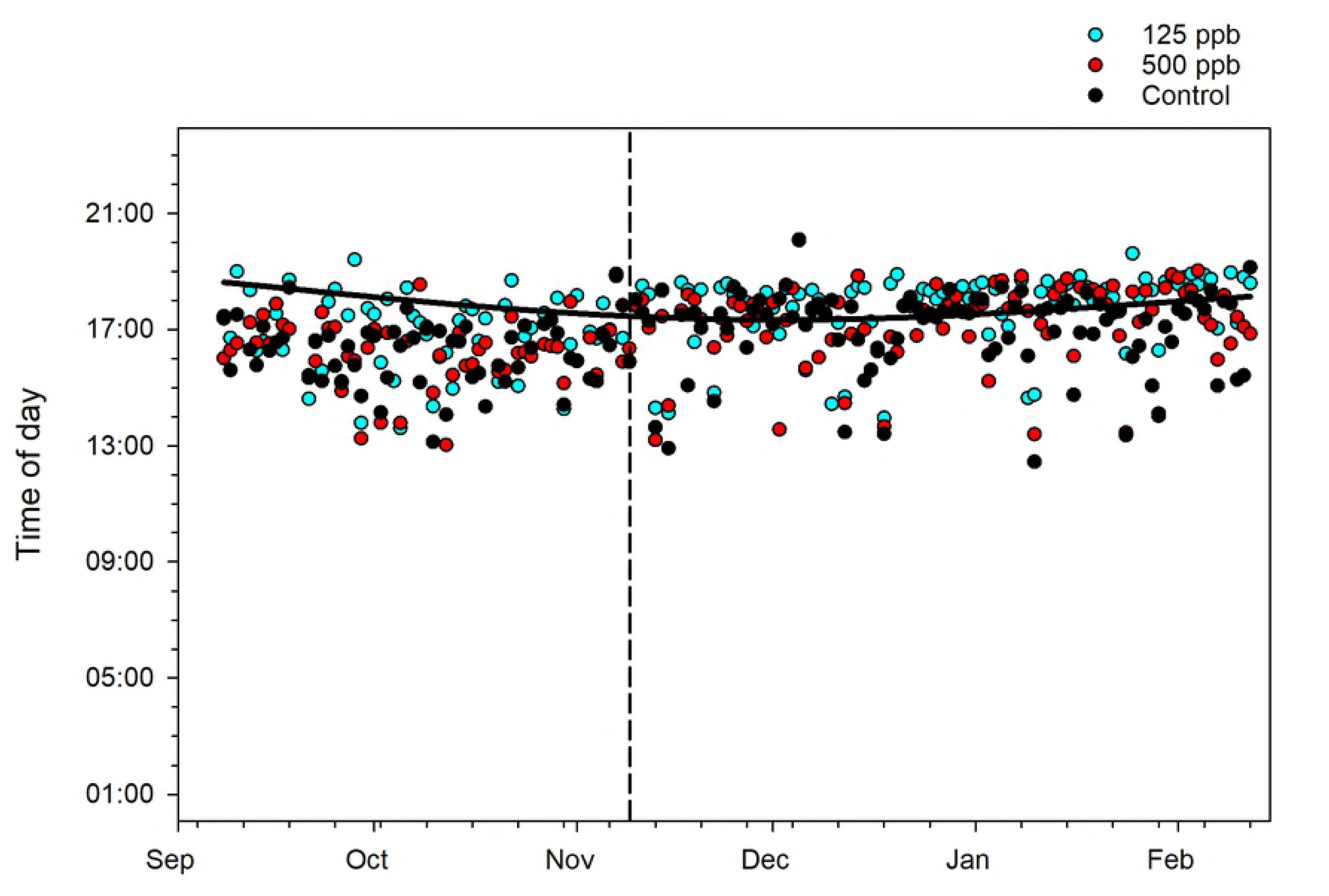
Break points associated with the end of the daily active period (dusk) for piecewise regression curves on within-day hive weight changes for the Fall 2017 experiment. Dashed black vertical line shows the end of the treatment period. Solid black horizontal line shows calculated sunset time.

Slopes of the segments associated with forager departure after the dawn break point were significantly affected by treatment (Fig 3, S1 Table). Post hoc contrasts showed that slopes in the 500 ppb treatment were significantly shallower (lower in absolute value) than those in the Control treatment (P=0.0043), indicating a lower rate of forager departure. Neither the night segment slopes nor the dawn break point were significantly affected by treatment (P=0.51 and 0.34, respectively).

**Figure 3.**
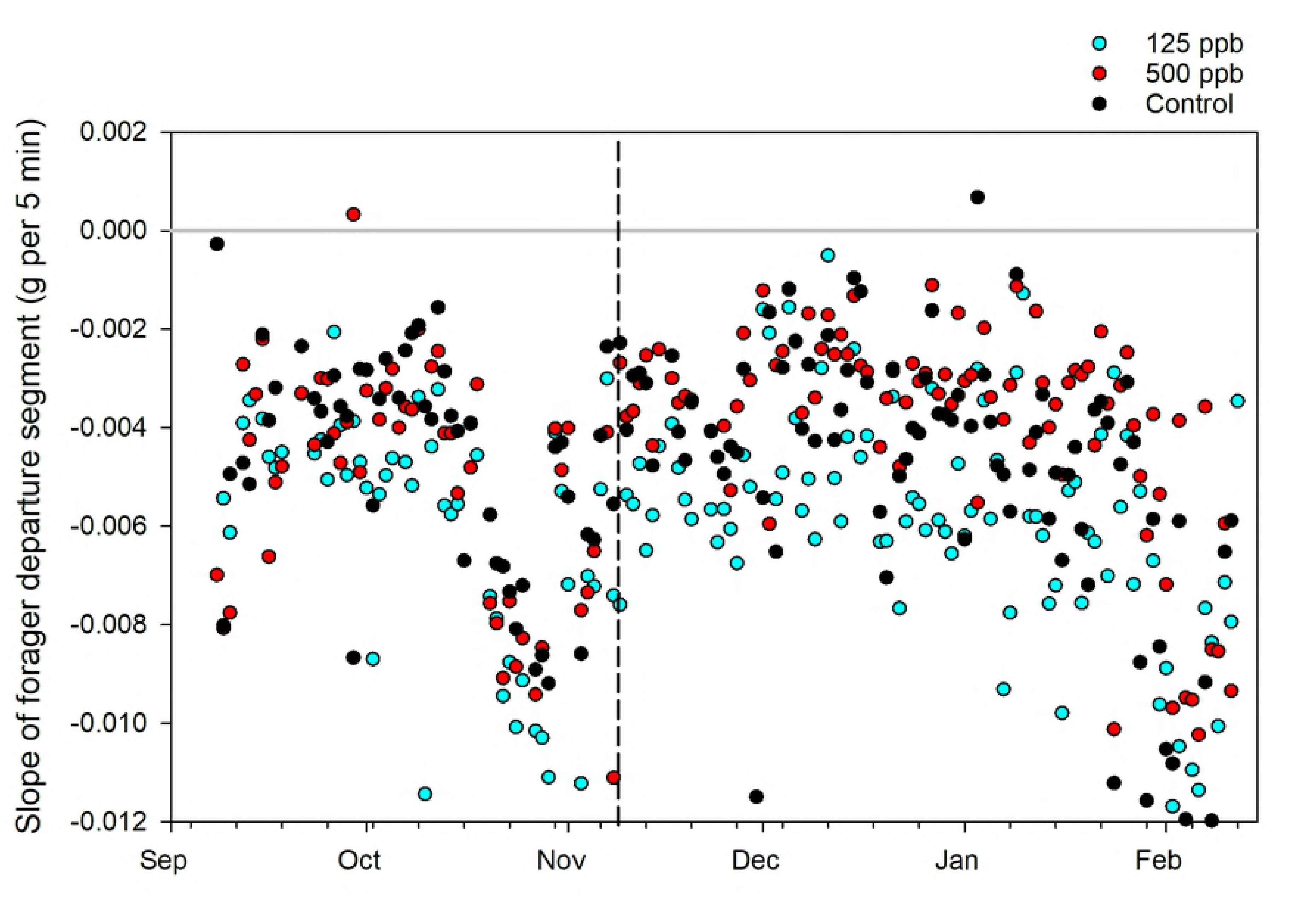
Segment slopes associated with departing foragers for the Fall 2017 experiment. Dashed black vertical line shows the end of the treatment period. Solid gray horizon line shows slope=0.

Average piecewise regression curves were calculated for each treatment group by averaging slope and break point values across all hives and sample days (Fig 4). Because the post-treatment data were collected in the late fall and winter, with few foraging opportunities, all hives lost weight as the bees consumed food stores. Average slopes at night tended to be positive, probably due to higher ambient relative humidity (the woodenware of the hives can gain weight, as well as any open food cells within the hive). In both years, average curves for the 500 ppb treatment group were shallower than the control, probably indicating a lower foraging effort.

**Figure 4.**
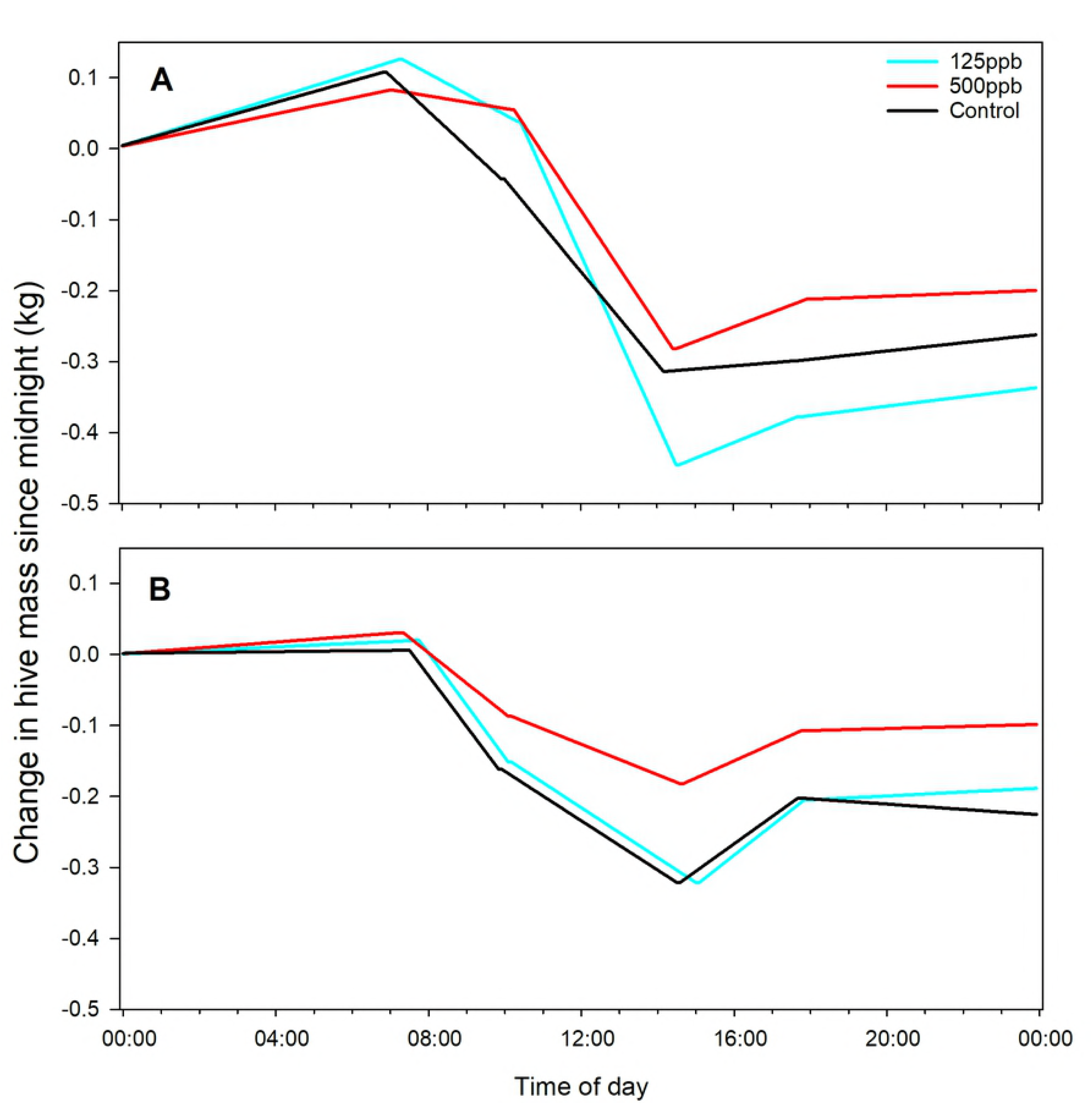
Piecewise regression curves calculated using average slope and break point values calculated from all colonies within each treatment group and across all days from the end of treatment until the final hive assessment. A) Fall 2016 experiment; B) Fall 2017 experiment. Graphs have the same scales.

Treatment groups within the same time period with no letters in common are significantly different at α=0.05 with a Bonferroni comparison for multiple groups. Bold indicates which group or groups had the highest value within each time period and response variable group. Dashes indicate no significant main effect.

### 7. Hive weight for Spring 2018 experiment

Hive weight consistently decreased during the fall experiments, but average hive weight in all treatment groups increased every day from the end of treatment on 20 April until 6 May. On 7 May average hive in each treatment group started to decrease, indicating the end of a nectar flow, and most hives lost weight after that day until the end of the experiment on 23 May. Within-day hive weight patterns differ depending on whether there is a nectar flow [25] so those two periods (20 April-6 May and 7-23 May) were considered separately. During the nectar flow, only the departing slopes were significantly different among treatments (S2 Table, Table 2). Average slope in the 500 ppb treatment group was significantly higher than the slopes for either of the other treatment groups. In this case, while average r^2^ ±s.e. values for model fit were high: 0.96±0.03, 0.97±0.03, and 0.91±0.06 for the 125 ppb, 500 ppb, and Control treatment groups, respectively, visual inspection of the data showed that breakpoints near dawn were not being detected by the algorithm, causing inaccuracies in other segment parameters such as slope values. Increasing the number of break points from 4 to 5 did not change the overall goodness of fit (average r^2^ =0.92, 0.97 and 0.96, respectively) and likewise did not improve detection of the break points (Fig. 5). These results, therefore, should be subject to future verification. No parameters were significantly affected by treatment during the 16 d after the end of the nectar flow.

**Table 2.** Results of post hoc comparisons for three response variables with significant treatment effects for three field experiments on the effects of sublethal methoxyfenozide exposure to honey bee colonies conducted in southern Arizona.

| Year | Concentration | Response variables |  |  |
| --- | --- | --- | --- | --- |
|  |  | Dusk break point | Departing slopes | Temperature amplitude |
| Fall 2016 | 500 ppb | **a** | **a** | **a** |
| & | 125 ppb | **b** | ab | ab |
| Fall 2017 | 0 ppb | c | b | b |
| Spring 2018 | 500 ppb | - | **a** | - |
|  | 125 ppb | - | ab | - |
|  | 0 ppb | - | b | - |

**Figure 5.**
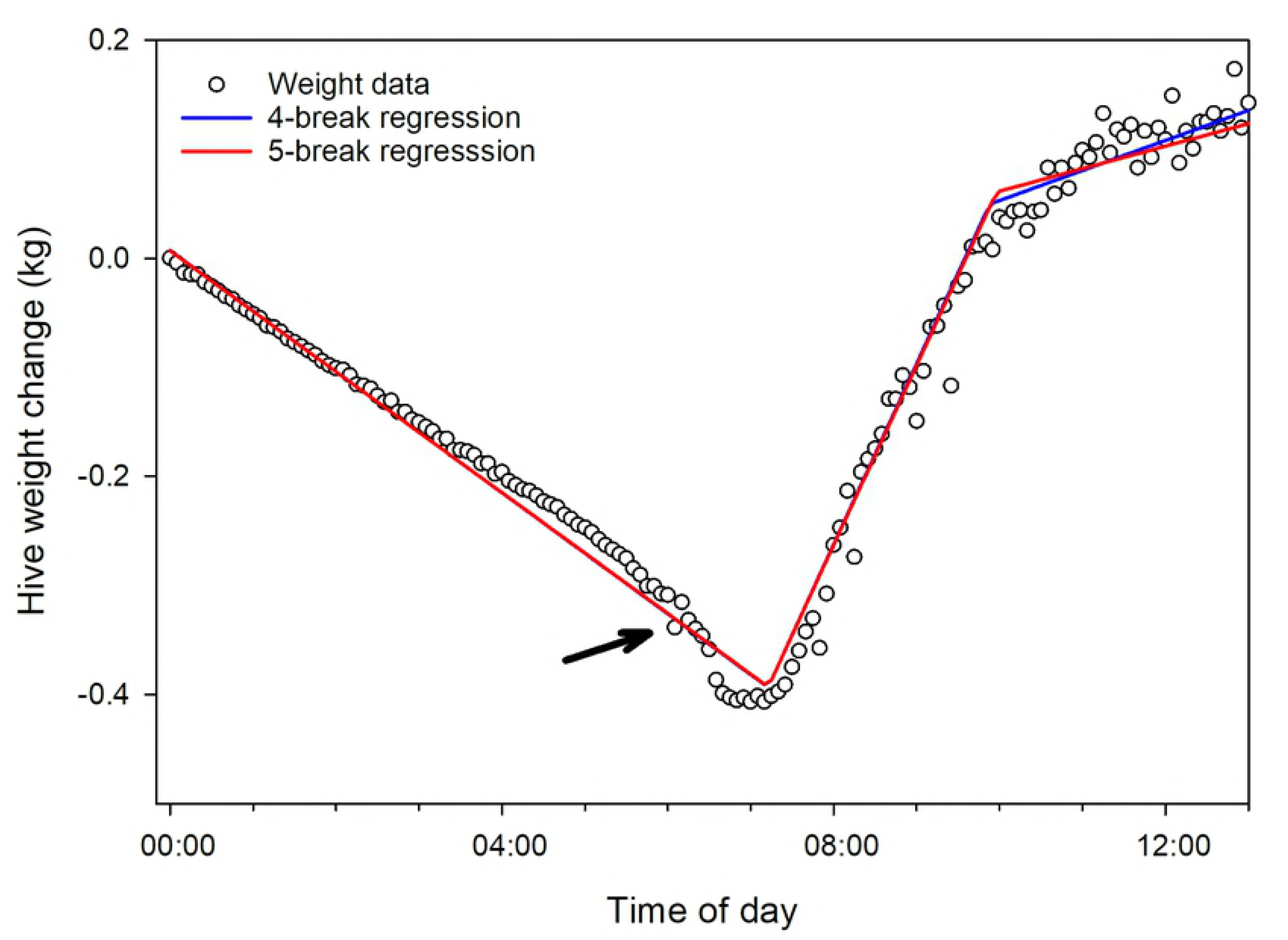
Example of an unsatisfactory curve fit. Raw data, collected on 29 April 2018 from a hive in the 500 ppb treatment group, shown with two piecewise regression curve fits: a curve with 4 break points and a curve with 5 break points. Arrow indicates expected “dawn” (1^st^) break point. Note that the 2^nd^ segment, typically associated with forager departure and therefore negative, is in this case positive.

### 8. Hive temperature for Fall 2016 and Fall 2017 experiments

Internal hive temperature was considered with respect to two response variables: 1) average daily temperature; and 2) average temperature variability measured as amplitudes of curves fit to 3-day datasets. Hive temperature has been positively correlated with total adult bee mass, so pre-treatment total adult bee mass was included as a covariate in all analyses. No significant treatment effects were observed either with respect to average temperature or to temperature amplitudes when all post treatment data (from end of treatment until final hive assessment), but the low P values (0.06 and 0.07, respectively) suggested there may be trends to explore by sharpening the focus of the analysis. Given that hive temperature is a function of both the bee colony and external ambient conditions, treatment effects may be more likely to be observed when a colony is challenged to manage its temperature. At the beginning of November in both 2016 and 2017 average ambient daily temperatures at the study site were about 21.7 to 22.8°C. Thirty days later, however, average ambient daily temperatures had dropped to 9.4-12.8°C. Thus, ambient temperatures 30 d after the end of treatment were considered more challenging to the bee colonies and thus more likely to show an effect. Considering only the data from 30 d after the end of treatment until the final hive assessment, treatment effects on average temperature remained not significant (P=0.06) but significant treatment effects were observed with respect to temperature amplitudes (variability) (S3 Table, Table 2, Fig 6). Temperature amplitudes were lower in the Control treatment group (about 1.72°C lower in the Fall 2016 experiment and 2.40°C in the Fall 2017 experiment) than the 500 ppb treatment group (about 2.55°C in the Fall 2016 experiment and 3.64°C in the Fall 2017 experiment). Neither the Control group nor the 500 ppb group was different from the 125 ppb group in either experiment.

**Figure 6.**
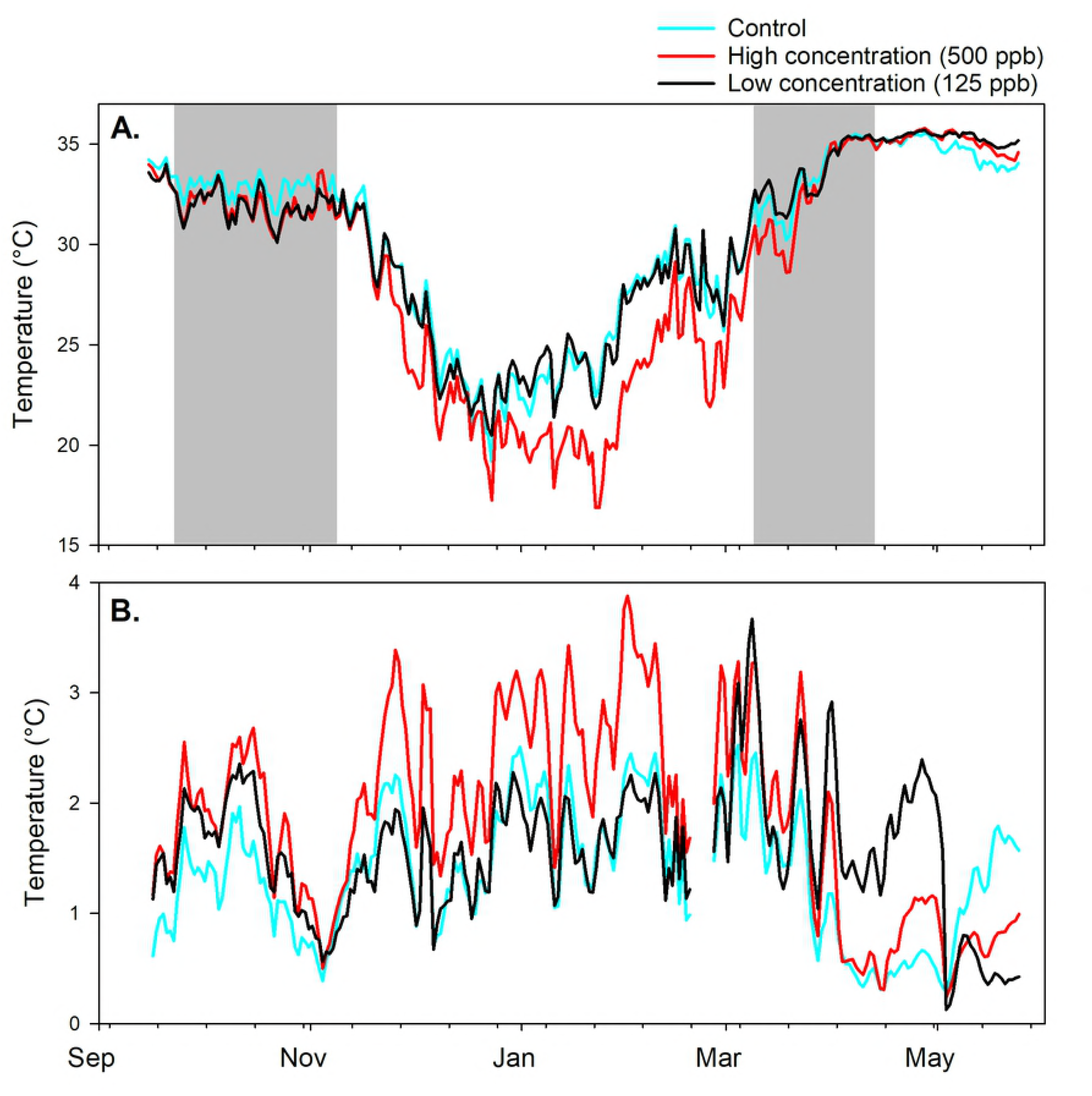
Internal hive temperatures. A) Average temperature for the Fall 2017 experiment; B) Average amplitudes of sine curves fit to internal hive temperature data (temperature variability). Gray zones in A) show the periods during which experimental treatments were applied.

### 9. Hive temperature data for Spring 2018 experiment

Neither average internal hive temperatures nor their daily variability were different among treatments (P=0.22 and 0.32, respectively) when the February adult bee mass was used as a covariate. February adult bee mass was significantly correlated with both temperature variables.

## Discussion

Methoxyfenozide is preferred as a pesticide for many crops pollinated by honey bees. One important question regarding sublethal exposure of methoxyfenozide is how the treatment effects manifest themselves, such as by reducing brood production or adult bee survivorship, changing colony behavior, or affecting the growth or physiology of individual bees. Methoxyfenozide, with a reported acute oral toxicity of more than 5.0 g per kg for humans, is considered mildly toxic for many vertebrates and crustaceans; with respect to honey bees, it has been reported as “moderately toxic” [9] and “relatively nontoxic” [32]. Most studies on sublethal pesticide exposure rely on invasive hive assessments to estimate adult bee and brood populations. To avoid antagonizing the bees or losing the queen, these inspections are typically carried out about every 4-6 weeks, as was done here. No effects of methoxyfenozide were observed with respect to hive assessment data (total adult bee mass or brood surface area) in this study. These data are important, but they provide little information on daily or hourly changes in colony-level behavior such as foraging and thermoregulation. In this study we monitored colony behavior using hive scales, to observe hourly weight changes associated with, for example, foraging activity [26], and using temperature sensors to measure themoregulation. Both types of sensors have been successfully used to detect the effects of sublethal exposure of honey bees to a neonicotinoid, imidacloprid [24].

Hive weight data, after being detrended by removing the value at midnight from subsequent values for the next 24 h, have common patterns [26]. For example, from midnight until the start of the active period, usually just after dawn, bee flight activity is minimal and hive weight changes largely involve water gain or loss depending on the amount of open nectar and ambient relative humidity. After dawn, weight changes usually become sharply negative, as the daily active period for the colony begins and foragers and other bees leave the hive. Whether hives gain or lose weight during the day depends largely on factors such as the success of the foraging bees. In the late afternoon, hives tend to gain weight as flying bees return to the colony; their return is usually complete about dusk, marking the end of the active period. After that point, hive weight changes are once again largely due to the amounts of hydrophilic materials present and to internal and ambient humidity. Hive scale data revealed treatment effects with respect to the slope of the segment associated with forager departure, and the dusk break point. Because these parameters resulted from the fit of single piecewise regression curves, they should not be considered entirely independent from each other but rather reflect fundamental differences in overall daily curve shape. Average curve shapes differed from year to year, showing the effects of yearly variability in temperature and rainfall, but clear differences in curve patterns were also evident within each year. Colonies fed 500 ppb methoxyfenozide in the fall in general had shallower curves, indicating lower activity than colonies in the Control group, and the 500 ppb colonies also had delays in the dusk break point of 17-21 minutes on average, consistent between the two fall experiments. Analyses of break points other than dawn, or of slopes other than the initial forager slope, were not included here because the meaning of any differences was not clear. Although the curves usually have similar shapes among colonies and over time, not all parts of the curves have a clear interpretation. This is particularly true during a forage dearth, when forage-related environmental signals are absent.

The Fall 2017 field experiment was continued through the following spring, in order to observe longer-term effects of methoxyfenozide exposure. The spring environment differed from that of the fall in two crucial respects: 1) rising temperatures, longer days and increasingly available forage promote colony growth rather than stasis or decrease as observed in the fall; and 2) a nectar flow was under way during the post-treatment period in the spring but not in either fall experiment. That the colonies grew rapidly in the spring in spite of reduced pollen patty consumption suggests that alternative food sources played a large role. Interestingly, few significant colony-level effects were observed in the spring. Three colonies died after the first hive assessment in the spring, one in the 500 ppb treatment and the other two in the 125 ppb treatment. As in the fall, hive assessment data were not significantly different among groups. Continuous weight data segment slopes associated with departing bees were significantly affected, with the 500 ppb hives showing higher slope values (indicating fewer departing bees) than other treatment groups, but this result did not conform with visual inspection of the data, reducing our confidence in that result. Regression models with 5 breaks, rather than 4, were fit to the data but the quality of the fit was not improved and merit further analysis. While the expectation was that hives may be more sensitive to IGR in the spring, IGR effects may have been diluted by the pollen and nectar flows (see [23]) or the bees, either on the individual or colony level, were more effective at detoxification.

Hive internal temperature has been correlated with adult bee mass and brood levels [23]. While adult bee and brood levels were not significantly affected by treatment, internal hive temperature was significantly more variable in the 500 ppb treatment group when November data were excluded (mild temperatures in November would have resulted in lower variability values in any case; removing those data put the focus on December through the final hive assessment in February). It may be that the temperature data were more sensitive to brood levels than the inspection data, if only because of the larger amount of temperature data, and as a consequence the temperature data revealed smaller differences.

Methoxyfenozide concentration was measured in wax samples during the Fall 2016 experiment, and none was detected in any sample. Wax was sampled to determine if the lipophilic nature of the compound facilitated its spread throughout the hive and that none was detected suggests that such spreading occurs at very small quantities, if at all. For the Fall 2017 and Spring 2018 experiments, bee bread was sampled. Low concentrations of methoxyfenozide were detected in bee bread from the 500 ppb treatment group and none in the other groups. The concentrations were apparently stable over time, from the end of the fall treatment in November until the end of the spring treatment the following April. Treatment patties were also sampled for methoxyfenozide concentration. Observed concentrations in the treatment patties were 17.6% lower than calculated concentrations in the 125 ppb treatment and 60.2% lower than calculated concentrations in the 500 ppb treatment. Disparities at about that magnitude between calculated and observed concentrations of pesticides mixed in pollen patties have been reported elsewhere [34]. Such disparities may be due to several factors, including insufficient mixing in a heterogeneous material (pollen patty), or a breakdown of the compound due to chemical reactions or biological activity in the patty environment. Using the observed concentration values, the bee bread results suggest that 1-2.5% of the bee bread sampled was treatment patty.

While significant effects of methoxyfenozide exposure on colony behavior were observed, no differences were detected with respect to hive assessment data (adult bee mass and brood surface area) or on the level of the individual bee (gland size, NEB dry weight and head mass of nurse bees). The kinds of colony-level behaviors that were measured, i.e. foraging activity and thermoregulation, can be considered functions of either colony size and age structure, or individual adult bee behavior, or both. Greater foraging activity by one colony compared to another may result from a larger adult bee population with a similar proportion of foragers or from a similar adult bee population with a higher proportion of foragers. Reduced variability in internal hive temperatures may result from the better insulation properties of a tighter bee cluster, from more heat, on average, per bee, or simply from more bees. Although brood levels were not significantly different, the 500 ppb treatment group ranked last for each post-treatment hive assessment in both fall experiments, suggesting that thermoregulation differences were likely (but not definitively) linked to brood levels. The significance of the observed delay in the end of foraging activity (“dusk”) is not clear, but longer-term effects of sublethal pesticide exposure are not always well understood. Six of the 36 colonies involved in this study died and none was from the control group. Further work is needed to link observed changes in colony behavior to longer-term effects on colony performance and survivorship.

## Conclusions

- Exposure of honey bee colonies to methoxyfenozide in supplement patty at a concentration of 500 ppb in the fall reduced the colony foraging population, delayed the end of the daily activity period by 17-21 minutes, and was associated with higher internal hive temperature variability (poorer thermoregulation).
- Exposure to the colonies did not have a measurable affect on the total adult bee mass, the amount of brood, average newly-emerged bee body mass or head weight, and caged bees fed 1000 ppb methoxyfenozide in sugar syrup showed no differences in hypopharyngeal gland size.
- Colonies treated for a 2^nd^ consecutive time the following spring showed fewer differences than in the fall.
- Continuous weight and temperature monitoring methods showed significant effects on colony-level behavior whereas periodic colony assessments and sampling did not show effects.

## Acknowledgements

The authors would like to warmly thank M. Heitlinger at the Santa Rita Experimental Range and M. Mcclaran at the University of Arizona for providing field sites for the work. In addition, the authors would like to thank S. Cook, V. Ricigliano and to two anonymous reviewers for their helpful suggestions on the paper.

---

**S1 Table.** Effects of treatment on dawn break point for piecewise regressions fit to continuous weight data for the Fall 2016 and Fall 2017 field experiments (PDF).

**S2 Table**. Effects of methoxyfenozide exposure on departing slopes for piecewise regressions fit to continuous weight data for the Spring 2018 field experiment (PDF).

**S3 Table.** The effects of methoxyfenozide exposure on log amplitudes of sine waves fit to detrended continuous temperature data (daily internal hive temperature variation) for the Fall 2016 and Fall 2017 field experiments 30 d after the end of treatment until the final assessment (PDF).

**S1 File**. Experimental data (XLSX).

